# Computational study on Strontium ion modified Fibronectin-Hydroxyapatite interaction

**DOI:** 10.1101/2021.09.23.461618

**Authors:** Subhadip Basu, Bikramjit Basu, Prabal K Maiti

## Abstract

Protein adsorption is the first key step in cell-material interaction. The initial phase of such adsorption process can only be probed using modelling approaches like molecular dynamics (MD) simulation. Despite a large number of studies on the adsorption behaviour of proteins on different biomaterials including hydroxyapatite (HA); little attention has been paid towards quantitative assessment of the effects of various physicochemical influencers like surface modification, pH, and ionic strength. Among these factors, surface modification through isomorphic substitution of foreign ions inside the apatite structure is of particular interest in the context of protein-HA interaction as it is widely used to tailor the biological response of HA. Given this background, we present here the molecular-level understanding of fibronectin (FN) adsorption mechanism and kinetics on Sr^2+^-doped HA (001) surface, at 300K by means of all-atom molecular dynamics simulation. Electrostatic interaction involved in adsorption of FN on HA was found to be significantly modified in presence of Sr^2+^ doping in apatite lattice. In harmony with the published experimental observations, the Sr-doped surface was found to better support FN adhesion compared to pure HA, with 10 mol% Sr-doped HA exhibiting best FN adsorption. Sr^2+^ ions also influence the stability of the secondary structure of FN, as observed from the root mean square deviation (RMSD) and root mean square fluctuation (RMSF) analysis. The presence of Sr^2+^ enhances the flexibility of specific residues (residue no. 20-44, 74-88) of the FN module. Rupture forces to disentangle FN from the biomaterials surface, obtained from steered molecular dynamics (SMD) simulations, were found to corroborate well with the results of equilibrium MD simulations. One particular observation is that, the availability of RGD motif for the interaction with cell surface receptor integrin is not significantly influenced by Sr^2+^ substitution. Summarizing, the present work establishes a quantitative foundation towards the molecular basis of the earlier experimentally validated better cytocompatibility of Sr-doped HA.

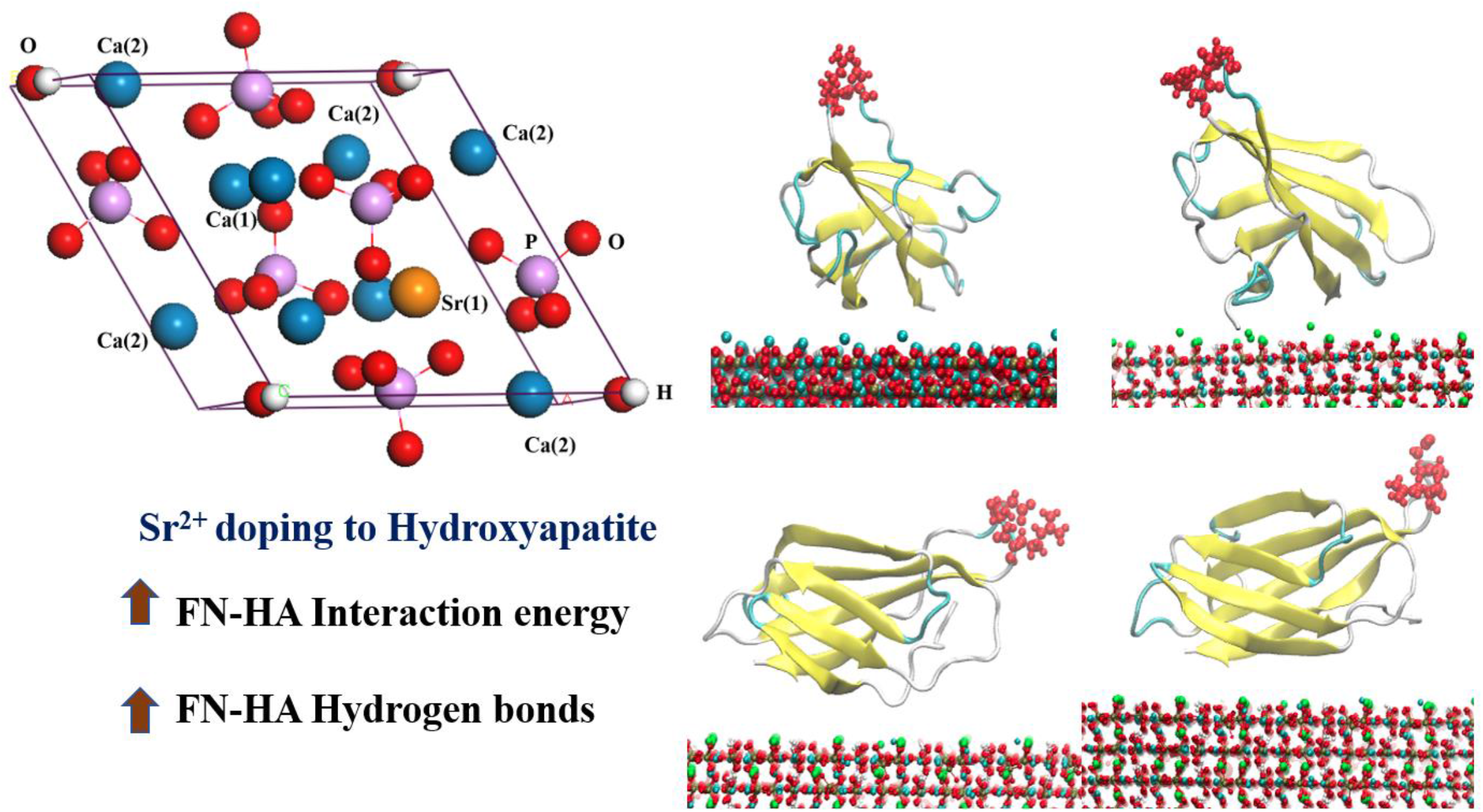

## 1. Introduction

Protein adsorption on materials surface is of utmost importance in the field of biomaterial science because of its central role in the tissue-material interaction.^1^ For example, adsorbed proteins mediate the cell adhesion on the biomaterials surface by taking part in the formation of focal adhesion complexes. Such interactions strongly depend on the final conformations/orientations of the adsorbed proteins.^2^ Moreover, adsorbed proteins play a major role in triggering the inflammatory response.^3^ Bacterial adhesion on a surface is also regulated by the nature of the adsorbed proteins, and it is known that bacteria exhibit different affinities towards different proteins.^4^ The protein adhesion on different substrates is also used by many living organisms, like sea urchins and mussels to adapt to the surrounding environment.^5^ Because of such widespread significances and multi-faced role played by protein-inorganic surface interaction, it is important to develop a deep understanding using computational analysis and experimental tools to get more insights into such interactions at multiple length scales.

Various experimental techniques like atomic force microscopy (AFM) and time of flight secondary mass spectroscopy (ToF SIMS) have been previously employed to probe the protein-material interplay with considerable success.^6, 7, 8^ However, with the development of computational techniques and computational power, different modelling approaches have been progressively used for quite some time to unravel the mechanisms of protein-materials interactions which cannot be probed by experimental means. The interaction of proteins, peptides, and amino acids with different kinds of surfaces like Au,^9, 10, 11^ SiO_2_,^12, 13, 14, 15, 16^ TiO_2_,^17, 18^ graphite,^17, 18^, polymers,^19^ self-assembled monolayers (SAMs)^20, 21^ have been studied using molecular dynamics (MD) simulation and density functional theory (DFT). This has helped in unveiling molecular/electronic level insights into the nature of the interplay between the surface and proteins, together with the information regarding the structural changes of proteins during adsorption phenomena.

One material that is of particular interest in the context of protein adsorption is hydroxyapatite (HA; Ca_10_(PO_4_)_6_(OH)_2_) because of its resemblance with natural bone mineral.^22^ Several researchers probed into the adsorption of various proteins on different surfaces of HA crystal lattice.^23, 24, 25, 26, 27^ It is evident from these studies that the adsorption of proteins on HA surface is mainly driven by electrostatic interaction among the oppositely charged groups together with the formation of H-bonds/water bridged H-bonds. Such phenomenon are particularly recorded in the cases of adhesion of bone morphogenic protein-2 (BMP-2) and BMP-7 on different HA surfaces.^28, 27^ Similar findings were reported for another important extracellular matrix protein, fibronectin (FN).^29, 30, 31^ FN is a dimer protein consists of three types of FN repeats, namely FNI, FNII and FNIII.^32^ While the long-range coulombic attraction was established to the driving force behind the adhesion of FNIII_10_ module on HA (001) surface,^29^ Van der Waals (VDW) force driven weak adsorbed state was observed in case of FNIII_7-10_ module.^30^

Despite all the above-mentioned studies, the effects of different physicochemical factors like pH, presence of different ions, surface modification, external stimuli (thermal, electrical, or magnetic) on the protein adsorption kinetics on HA surface is hardly explored. Among these influencing factors, the effects of surface modification through doping of ions inside the apatite structure on the behaviour of adsorbed proteins are of particular interest because the method of isomorphic substitution is often used to overcome the limitations of pure HA for biomedical applications.^33^ Among different ion-doped HA, Sr^2+^-substituted HA is widely explored in the biomaterials community, because of the well-known beneficiary effects of Sr^2+^ on bone-cell growth.^34^ Sr supports bone regeneration by enhancing osteoblastic activity and suppressing osteoclastic activity.^35^ Sr^2+^ also promotes osteogenic differentiation by positively affecting CaSR and downstream signaling pathways.^36^ Better protein adhesion was found on Sr-doped HA surface compared to pure HA, when tested with bovine serum albumin (BSA) lysozyme, fibronectin, and vitronectin.^37, 38^ the molecular basis of such behaviour of Sr-doped HA is still unknown. In a recent study,^39^ adsorption behaviour of BMP-2 on Sr-doped HA has been investigated using MD simulation technique. They have reported an increase in the interaction energy and the number of contacts between protein and surface, in presence of Sr^2+. 39^ There is a necessity to carry out similar studies on other proteins, as adsorption kinetics is protein specific. Moreover, there is a recent development of force field parameters of Sr^2+^ ions inside apatite structure, which is expected to provide a better insight into the biological response of Sr-doped HA.^40^ Against this background, the present study aimed to explore the adsorption behaviour of FNIII_10_ module on Sr-doped HA (001) surface using MD and steered MD simulation. The influences of Sr^2+^ ions on the structural stability of FN have been quantitatively investigated in terms of RMSD and RMSF analysis of protein module. Dopant-dependent interaction between FN and material surface has also been thoroughly analyzed, together with the determination of force required to detach the protein from the surface. In the end, interdependence among the obtained results has been established.

## 2. Methods

### 2.1 Model of the protein

FN-III_10_ module was chosen as the model protein. The structure of FN-III_10_ was taken from Protein Data Bank (PDB id 1TTF). It consists of 94 residues with a molecular weight of 9.95 kDa.^41^ The net charge of this module is 0 together with a total dipole moment of 245.7 D.^42^ This particular module was selected for the present study because of the presence of the RGD motif in it.^41^

### 2.2 HA surface

The unit cell of HA features the following crystallographic parameters: *a=b=*9.432Å, *c=*, 6.881Å, *α=β=*90°, *γ=*120°. The Sr^2+^ ions were substituted in Ca(1) positions with doping percentages of 10,20 and 30 mol%, in corroboration with our previous experimental study.^43^ The force field parameters for HA were taken from INTERFACE force field.^44^ The parameters for Sr^2+^ ion were obtained from the existing literature.^40^ Throughout the article, xSrHA/xSr implies x mol% Sr doped HA.

### 2.3 Simulation details

Slabs featuring 12×10×4 unit cells of pure and different Sr-doped HA were built and used for the study. A periodic box of size 16.3004×10.1554×12.04 nm^3^ was made and. the FN module was placed ∼5Å above the (001) facet of the biomaterials surface (HA/Sr^2+^ doped HA)with preferred adsorption orientation.^30^The box is then solvated with TIP3P water model. The snapshot of the system is shown in Fig. 1. This particular surface has been chosen because it is well-known that, {100} planes of HA play an important role in interaction with biomolecules and the overall growth of HA crystal occurs in (001) direction during biomineralization of hard tissues.^45^ The energy minimization was performed in 10,000 steps using the steepest descent algorithm to remove close contacts between atoms. The system was then equilibrated at 300K and 1 bar using MD simulation in NVT and NPT ensemble for 1ns and 2ns, respectively. Modified Berendsen thermostat^46^ and Parrinello−Rahman barostat^47^ were employed for temperature and pressure regulation, respectively. Position restraints were applied on FN and material surface (HA/Sr^2+^ doped HA) during equilibration. The equilibrated system was then used for the production run (NPT) of 120ns with position restraints on biomaterials surface (HA/Sr^2+^ doped HA) only. The integration time step was 2fs throughout the simulation and the coordinate trajectory was recorded every 10ps for subsequent analysis. The coupling constant of barostat was kept at 0.5ps and 2ps during equilibration and production runs, respectively. We used 0.1ps coupling constant for the thermostat. LINCS algorithm^48^ was employed to constraint all bonds. Particle mesh Ewald method^49^ was used to calculate long range interaction. 10 Angstrom cut-off was used to compute non-bonded short-range interaction. All simulations were performed in GROMACS 5.1.4 package with AMBER99SB-ILDN force field for FN.^50, 51^ Calculations of different properties were carried out using GROMACS modules. VMD 1.9.3 software was used for visualization.^52^

**Fig 1:**
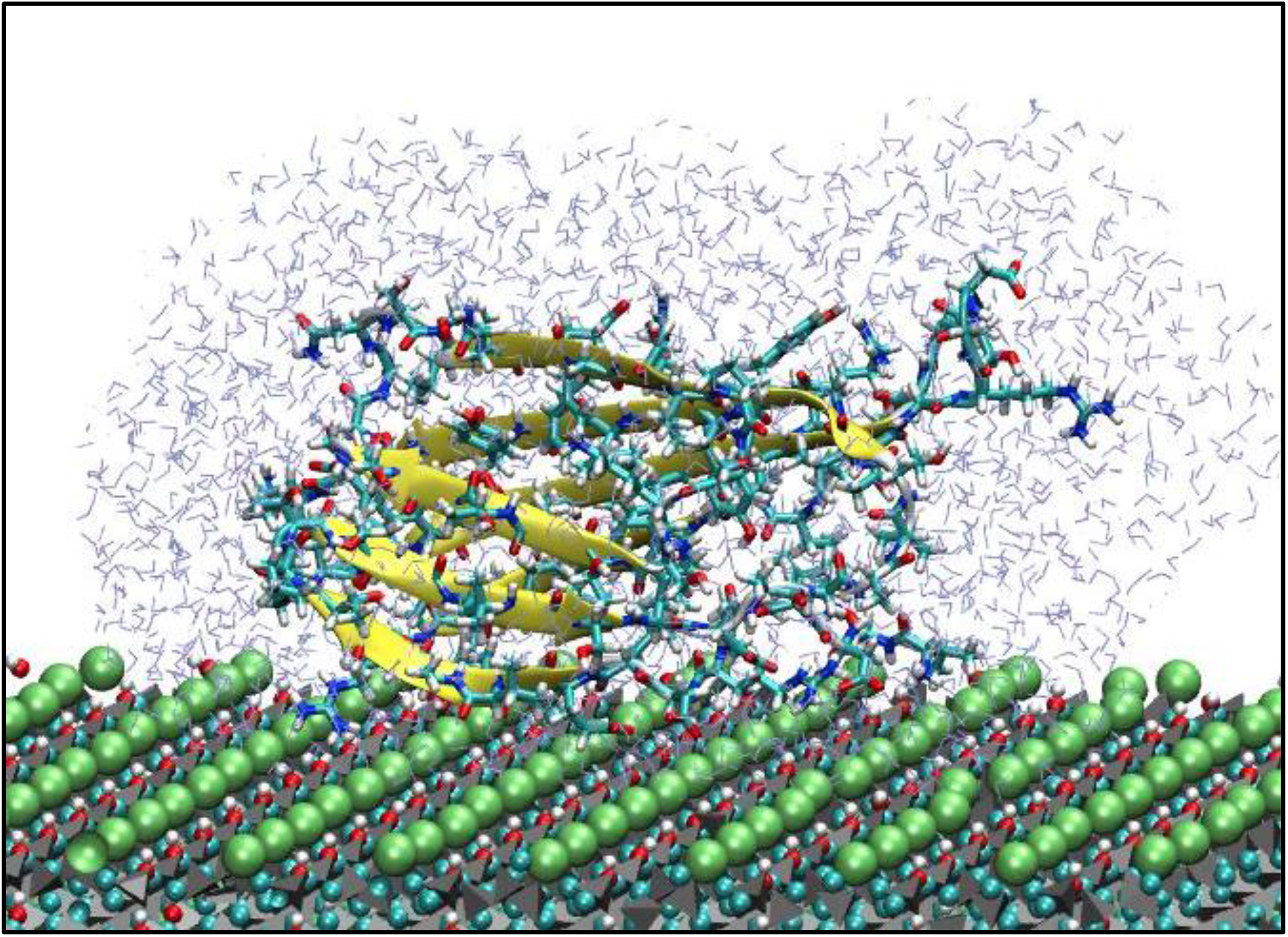
Initial configuration of FN-Sr-doped HA surface system. Only some of the Water molecules (light blue lines) are shown due to clarity. FN has been shown using the licorice representation with the colour scheme: Red: O, pale blue: C, white: H, blue: N. The secondary structure of FN has been represented with colour code: Yellow: β sheet, pale blue: turns, white: other residues. HA has been presented using CPK colouring scheme with atomic colour code: Red: O, pale blue: Ca, white: H, Green: Sr. Phosphate groups are presented using grey tetrahedrons.

Steered molecular dynamics (SMD) was performed for 500ps on the final configuration obtained after 120ns of equilibrium MD simulation. SMD was performed with a constant pulling velocity of 0.005 nm/ps. FN was being pulled away from the surface. During pulling, an elastic spring with a spring constant of 1500 kJ mol^−1^nm^−2^ was applied on FN module along the pulling direction, while the biomaterials surface was taken as static reference. Steering forces were recorded for every 1ps. Rupture force, defined as the highest force applied on the FN before detachment was calculated for each case together with the unbinding time.

## 3. Results

In the current section, the phenomenological behaviour of the FN module on different HA surfaces during the adsorption process will be penned.

### 3.1 Effects of Sr doping on Adsorption behaviour

In order to obtain a quantitative aspect of the adsorption behaviour of FN module during adsorption, the different modes of interaction between FN and biomaterials surface have been analyzed using GROMACS module. The representative results are presented in Fig. 2 and Table 1. It is evident from Fig. 2(a), that the long-range electrostatic interactions played the dominant role in the adsorption kinetics. In the case of undoped HA, the average interaction energy turned out to be - 523.9 ±101.9 kJ/mol. The interaction energy was close to ∼-450 kJ/mol for the first ∼40ns of the simulation and then it dropped to ∼-650 kJ/mol between 40ns to 80ns. After that, the interaction energy reached ∼-200 kJ/mol at ∼88ns. This recorded trend signifies a strong and weak adsorbed state of FN on HA surface.

**Table 1:** Average interaction energy between FN and surface, obtained from calculated trajectories

| Code of Biomaterials surface | Sr Doping (mol%) | Average coulombic interaction (kJ/mol) | Average non-bonded interaction (kJ/mol) |
| --- | --- | --- | --- |
| 0SrHA | 0 | $-523.9 \pm 101.9$ | $2.2 \pm 17.7$ |
| 10SrHA | 10 | $-794.2 \pm 85.6$ | $-17.5 \pm 19.1$ |
| 20SrHA | 20 | $-649.8 \pm 59.5$ | $17.7 \pm 17.8$ |
| 30SrHA | 30 | $-680.9 \pm 39.5$ | $2.8 \pm 17.4$ |

**Fig. 2:**
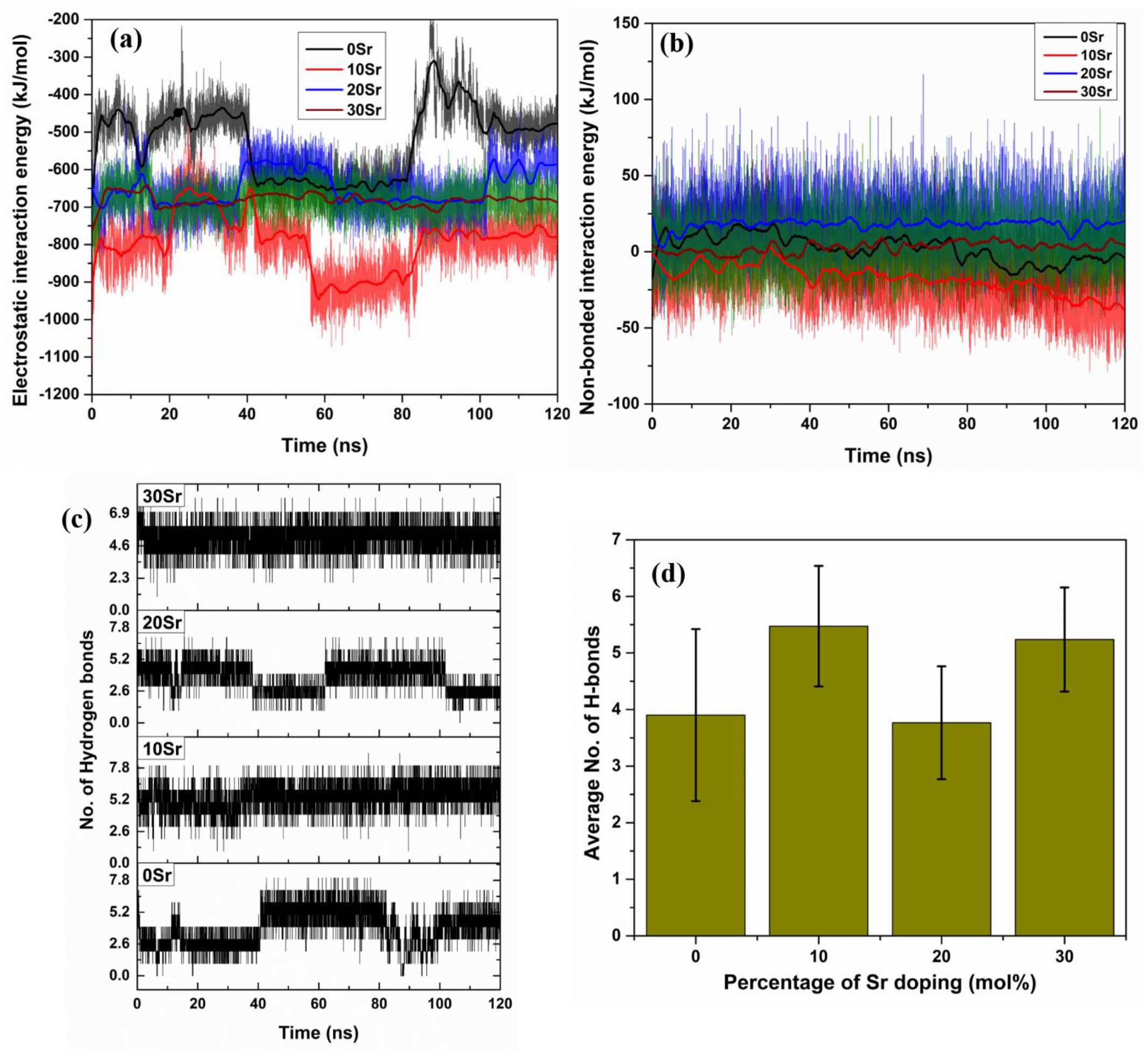
Doping of HA with Sr^2+^ profoundly affects different modes of interaction between FN and surfaces. (a) Long-range coulombic and (b) non-bonded interaction between FN and surfaces. (c) temporal evolution and (d) average number of Hydrogen bonds formed between surfaces and FN. Highest coulombic interaction and maximum number of H-bonds was found for 10SrHA surface. In (a) and (b), the legend colour codes represent the thick lines (generated using Savitzky–Golay filter).

The attractive coulombic interaction energy increased in the presence of Sr^2+^ ions, as seen in Fig 2(a). The maximum interaction energy estimated was in the case of 10 mol% Sr-doped HA surface (−794.2±85.6 kJ/mol). The interaction energy decreased with an increase of further Sr^2+^ content (Fig. 2(a)). However, the interaction energy was always higher for Sr-doped surfaces compared to undoped one (Fig. 2(a) and Table 1). This indicates that, the presence of Sr^2+^ ions promotes adsorption and 10 mol% Sr-doped HA (10SrHA) presents the thermodynamically most favourable surface for adsorption. Also, the major contribution towards electrostatic interaction energy comes from the interaction of FN with Sr^2+^ ions, probably because of its large ionic radius (Fig. S1 in supporting information). On the other hand, there is no significant change in the non-bonded LJ interaction energy (Fig. 2(b)). The magnitude of non-bonded LJ interaction energy was found to be much less compared to its electrostatic counterpart (Fig. 2(b)).

Adsorption is often influenced by the formation of hydrogen bonds between protein and the material surface. For the present system under study, the oxygen atoms of the hydroxyl groups of the surface act as H-bond donors and all oxygen atoms play the role of acceptors. On the other side, the oxygen atoms of the side chains of Ser, Thr and nitrogen atoms of the side chains of Lys, Arg together with the nitrogen atoms in the main chain peptide bonds of FN module can act as donors/acceptors. The donor-acceptor cut-off distance was 3.5Å. The temporal evolution of the number of H-bonds formed between FN and Sr-doped HA surface is depicted in Fig. 2(c). The average number of H-bonds was plotted in Fig. 2(d). The average number of H-bonds was higher for 10SrHA (5.5 ±1.1) and 30SrHA (5.2 ±0.9) surface than the undoped one (3.9±1.5), and was slightly low for 20SrHA (3.8±0.9). In corroboration with the previously observed results, this also indicates that, the adsorption of FN is most favoured on 10mol% Sr-doped HA.

Another important aspect in the context of adsorption phenomena is the protein-surface distance as a higher protein-surface distance indicates a weakly adsorbed or desorbed state.^31^ In the present study, the distance between the centre of mass (COM) of FN and surface has been computed together with the minimum distance between them (Fig 3(a)). The minimum distance is defined as the minimum of all the atomic pair distances, where one atom of the pair belongs to FN and other belongs to the surface. The COM-COM distance between FN and surface kept changing up to ∼40 ns of simulation time for 0SrHA, and after that, it remained constant at ∼2.95nm (Fig. 3(a)). For the doped surfaces, the distance remained nearly constant for the simulation time span (Fig. 3(a)), indicating a steady adsorbed state throughout. On the other side, the minimum distance between FN and surface always remained ∼0.16 nm, irrespective of the doping status (Fig. 3(b)). This is a signature of the affinity of FN towards the HA surface.

**Fig. 3:**
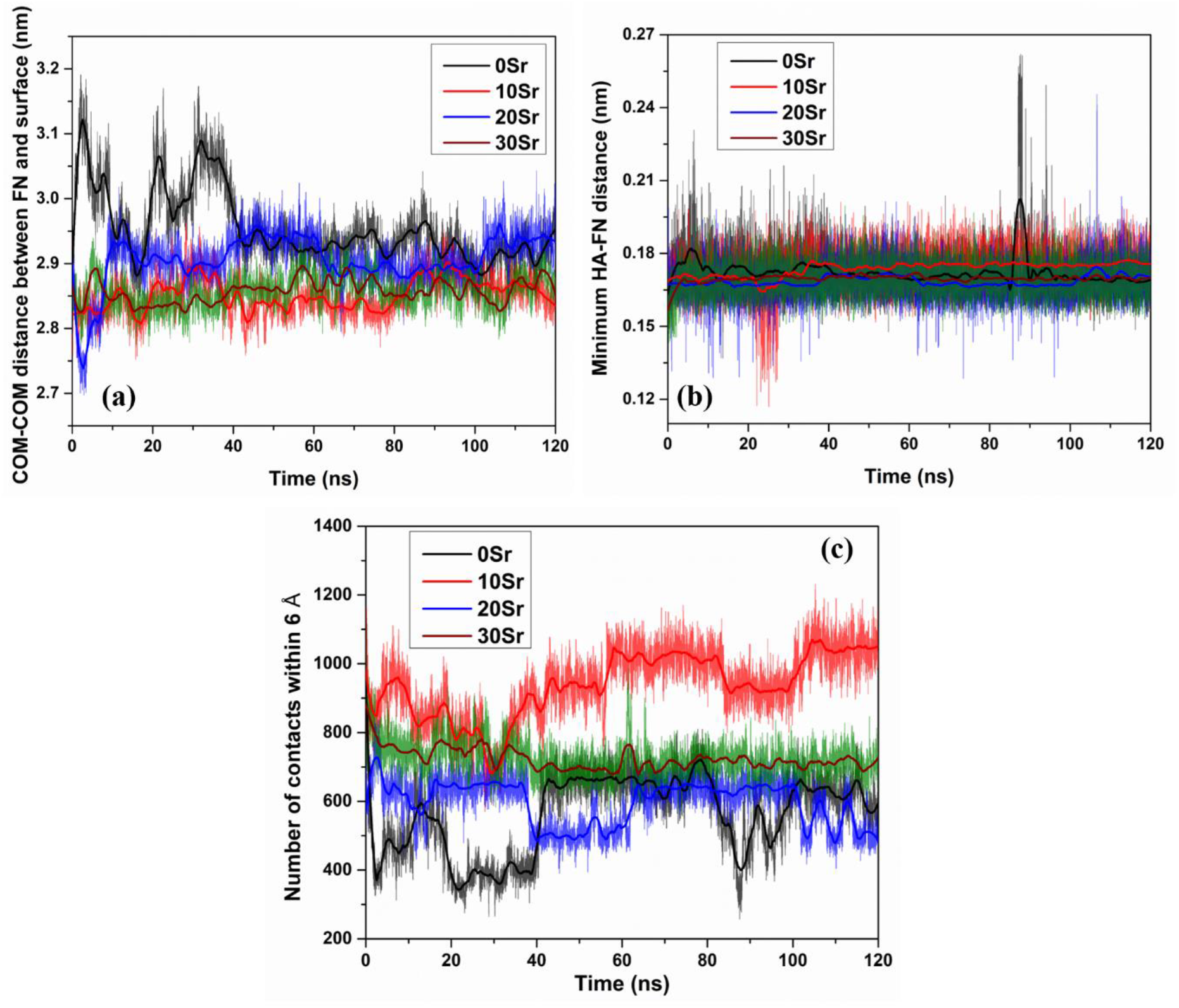
Different parameters, used for quantitative assessment of adsorbed states, is modulated on the surface of Sr-doped HA. (a) distance between centre of masses (COM) of FN and pure/Sr-doped HA substrate, together with (b) minimum distance, and (c) number of contacts between FN and biomaterials surface. In all graphs, the legend colour codes represent the thick lines (generated using Savitzky–Golay filter).

A quantitative measure of the adsorption is the number of contacts formed between FN and the biomaterials surface, which has been calculated, as shown in Fig. 3(c). It is noteworthy that a contact is defined when two atoms, one from the surface and other from FN, are within the cut-off distance of 6Å. For 0SrHA, the number of contacts fluctuates between ∼200 and ∼700. Such observation, in harmony with previously observed results, depicts a change in the adsorbed state of the protein (Fig.3(c)). Similar behaviour was found for 20SrHA surface (Fig. 3(c)). The highest number of contacts was found to be formed in the 10SrHA-FN system (Fig. 3(c)). This a symphonized proof of the fact that, the 10SrHA surface is the most favourable for the FN adsorption. For 30SrHA, the number of contacts was calculated to be higher than 0SrHA (Fig. 3(c)), which establishes the fact that 30SrHA is also suitable for adsorption of FN.

### 3.2 Adsorbing residues

It is now well-known that not all residues of FN interact equally with HA surface. Similar to our previous study,^31^ the charged residues were found to be interacting more strongly with the surface (and Fig. S5 and S6 in supporting information) and have been listed in Table 2. Positively charged residues interacted with the surface via NH_3_^+^ groups, while negatively charged residues interacted through COOH^-^ groups. The number of the interacting residues changed with surface modification, as seen in Table 2. Highest number of interacting residues was found on 10SrHA surface. The interaction energy of different residues, obtained using standard GROMACS module, has been plotted in Fig.4. For 0SrHA, Arg93 and Glu9 turned out to be the most interacting residues, followed by Arg6 (Fig.4(a)). On the other hand, the interaction between Asp7 and 0SrHA surface was repulsive in nature (Fig.4(a)). Hence, this particular residue did not favour the adsorption process.

**Table 2:** Adsorbing residues of FN on different Sr-doped/undoped HA surface.

| Code of Biomaterials surface | Sr Doping (mol%) | Adsorbed residues (within 5 Å) |
| --- | --- | --- |
| 0SrHA | 0 | Arg93, Arg6, Asp7, Glu9 |
| 10SrHA | 10 | Val1, Ser2, Arg6, Asp7, Glu9, Thr94, Arg93, Asn91, |
| 20SrHA | 20 | Arg6, Asp7, Glu9 |
| 30SrHA | 30 | Arg93, Glu9, Asp7, Arg6 |

**Fig. 4:**
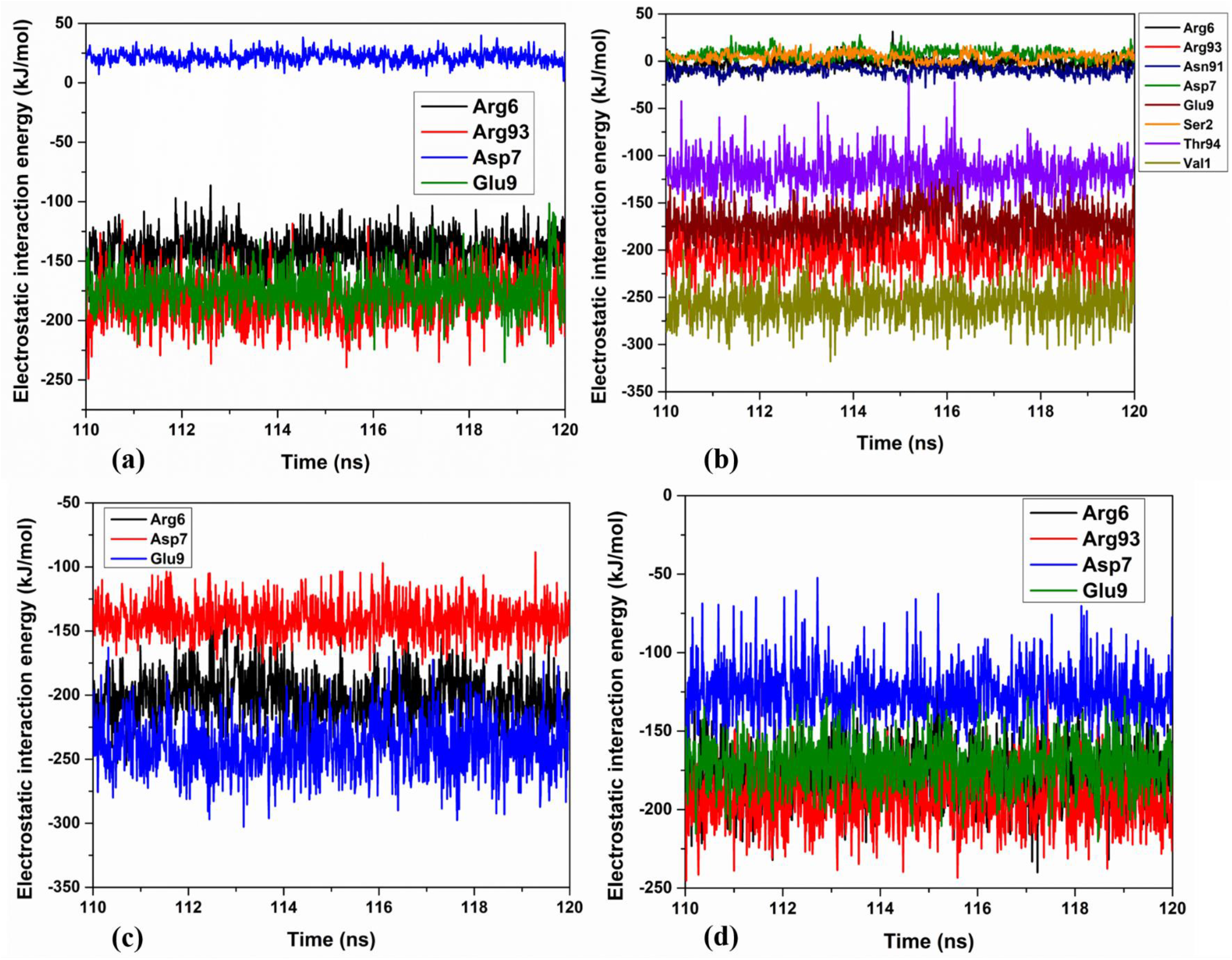
Number of adsorbing residues of FN together with the strength of interaction can be tailored by Sr^2+^ doping. Electrostatic interaction energies of different adsorbing residues of FN with (a)0SrHA, (b) 10SrHA, (c) 20SrHA and (d) 30SrHA.

In case of 10SrHA, Val1 was observed to interact most strongly with the surface (Fig. 4(b)). Arg93, Glu9, Thr94 also interacted strongly with the Sr-HA surface, promoting adsorption (Fig. 4(b)). On the other hand, Arg6, Asn91, Ser2, and asp7 interacted weakly with the Sr-HA surface and did not play much significant role in adsorption (Fig. 4(b)). The situation changed drastically with the further increase in the dopant content. As the number of interacting residues decreased with the increment in the number of Sr^2+^ ions, the total interaction energy decreased (Fig.4(c)). For 20SrHA, Arg6, Asp7, and Glu9 were the only interacting residues and all of these interactions promoted adsorption of FN (Fig. 4(c)). Similarly, Arg6, Arg93, Glu9 and Asp7 interacted with 30SrHA surface, to promote the adsorption phenomena (Fig. 4(d)). One interesting observation is that, the interaction of Asp7 with the surface grew stronger with an increase of the Sr^2+^ content. Moreover, Glu9 and Arg6 were the only two residues that interacted with the surfaces, irrespective of their composition (Table 2).

### 3.3 Sr-doping dependent conformational changes

The root mean square deviation (RMSD) analysis was carried out to probe the structural features of FN module during adsorption process. It has been noted that, the RMSD of the backbone of FN was highest in the case of 0SrHA surface and was found to be lower for all other surface modifications (10SrHA, 20SrHA and 30SrHA surfaces) (Fig. 5(a)). This implies that the conformation of FN module was more constrained on the Sr-doped surfaces, compared to their undoped counterpart. In order to figure out the effective residues involved in the maintenance of the FN structure, root mean square fluctuation (RMSF) of the C_α_ atoms of the residues were calculated and has been presented in Fig. 5(b). It was noticed that the RMSF of the residues exhibits similar patterns in all the cases and the major difference was observed in the areas consist of residue no. 36-44 (area1) and 74-88 (area2) followed by a region made of residue no. 20-36 (area3) (Fig.5(b)). In particular, ‘area1’ showed the highest fluctuations for 0SrHA whereas area2 exhibited the highest fluctuations for 30SrHA. This indicates towards the surface dependent modification of FN structure in order to accommodate in the modified environment. The low value of RMSF of the FN residues for 20SrHA surface can be correlated with the relatively stable value of RMSD throughout the simulation duration (Fig.5(a) and 5(b)) as lesser structural changes gave rise to lower fluctuations. Moreover, the high fluctuation recorded in ‘area2’ is associated with the change in the β-sheet structure on 30SrHA surface. A change in the % of β-sheet (From 0-∼55%) has been observed in that region (Fig S2 in supporting information).

**Fig. 5:**
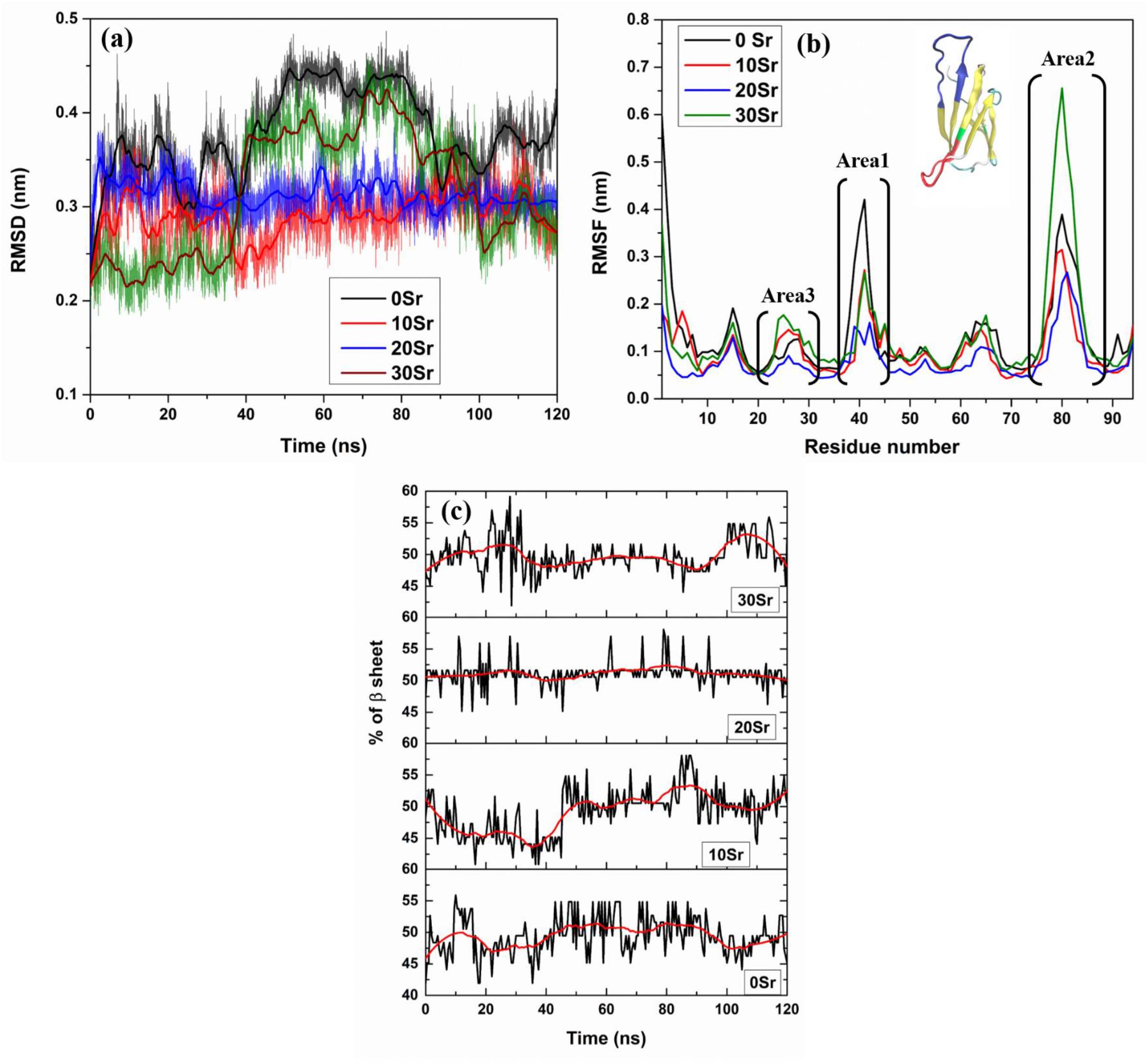
Sr^2+^ induces structural changes in FN, which in turn affect the adsorption kinetics. (a) time evolution of RMSD of backbone of FN, (b) RMSF of C_α_ atoms of FN residues, and (c) change in percentage of β-sheet with time. In (b), the most affected areas of FN by surface modification are marked and their positions is shown in FN module in inset with the colour code: red: area1, Blue: area2 and green: area3. In (a), the legend colour codes represent the thick lines (generated using Savitzky–Golay filter).

Besides this, the conformation of the protein has been altered in a manner dependent on Sr^2+^ content. In order to obtain a quantitative aspect, the temporal evolution of the percentage of β-sheet is plotted in Fig. 5(c). For 0SrHA, the percentage of β-sheet was found to be oscillating between ∼42% and ∼56% (Fig. 5(c)). The percentage of β-sheet remained almost stable at an average value of ∼51% between 48 ns and 84 ns (Fig. 5(c)). After that, the mean value dropped to ∼47% at ∼101 ns and raised slightly after that (Fig. 5(c)). For 10SrHA, the percentage of β-sheet first dropped from an average value of ∼50% to ∼44% at ∼36 ns and again rose to ∼50% at ∼52 ns and maintained a nearly steady-state for the rest of the simulation time (Fig. 5(c)). On the other hand, the percentage of β-sheet remained stable at around 50% value throughout the simulation window for 20SrHA surface (Fig. 5(c)). A significant change in the average value of % β-sheet has been recorded for 30SrHA between 92 ns and 120 ns (Fig. 5(c)).

In the context of the cell-material interaction, the position of RGD sequence is of utmost importance. If the RGD motif gets adsorbed on the surface, it will not be available for cell surface receptor integrin; and this is not favourable condition for cell adhesion on biomaterials surface. In the present study, the RGD motif was always found to be exposed in the solvent, irrespective of the biomaterial composition (Fig. 6). Hence, doping with Sr^2+^ ions does not affect cell-surface interaction.

**Fig 6:**
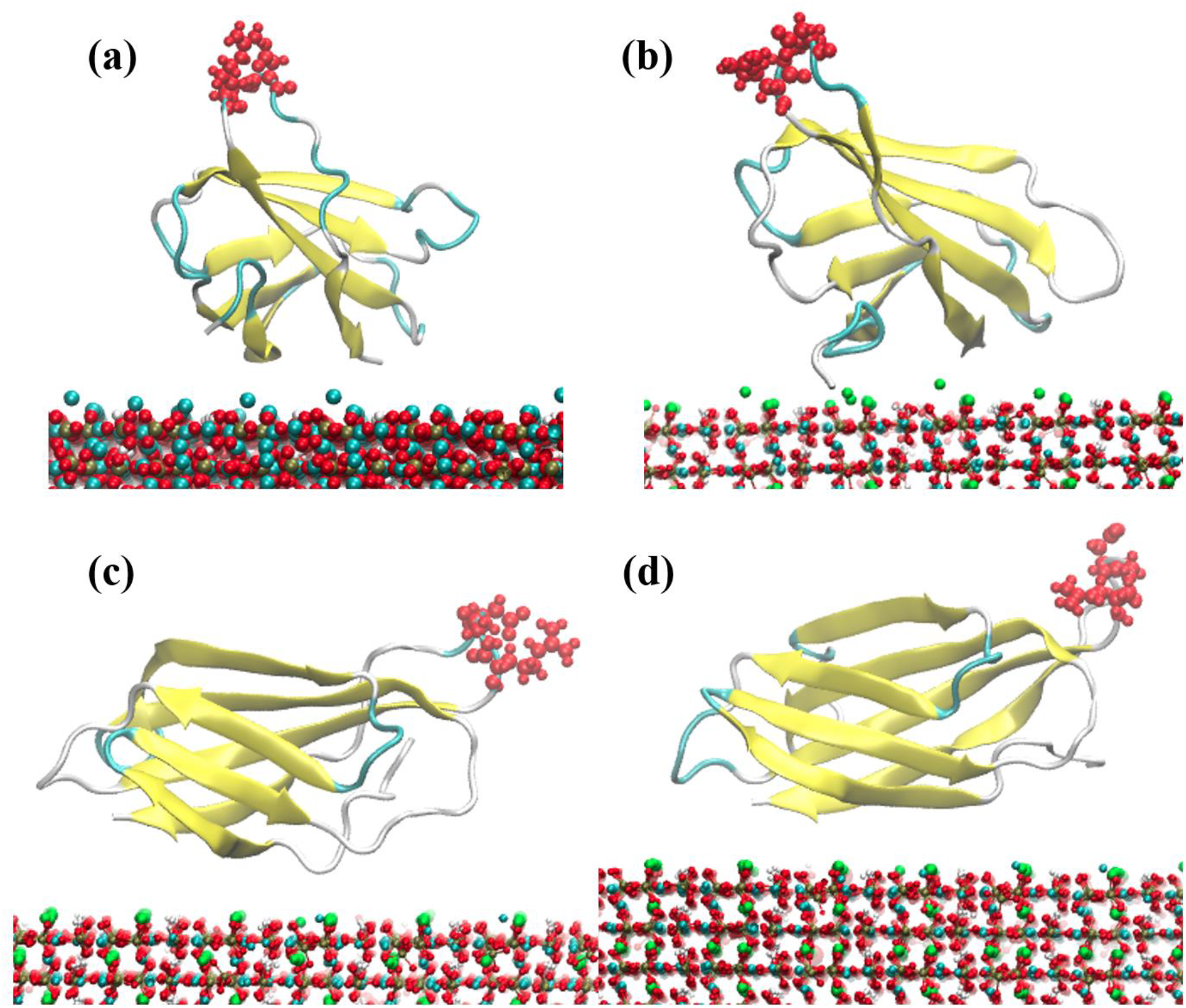
Cell-material interaction is not compromised in presence of Sr^2+^ ions inside HA crystal structure. Position of RGD motif (shown in red CPK spheres) on FN module adsorbed on (a) 0SrHA, (b) 10SrHA, (c) 20SrHA and (d) 30SrHA after 120ns of simulation. Water molecules are not shown due to clarity. The secondary structure of FN has been represented with colour code: Yellow: β sheet, pale blue: turns, white: other residues. HA has been presented using CPK colouring scheme with atomic colour code: Red: O, pale blue: Ca, white: H, golden yellow: P, Green: Sr.

### 3.4 Determination of unbinding time and rupture force

Typical unbinding curves of FN for each of the surface is presented in Fig. 7 together with the instantaneous snapshot of the system at the end of 500ps. Rupture forces and unbinding times of FN, adsorbed on different surfaces, have been listed in Table 3. From Fig. 7(a), it is clearly seen that the rupture force was highest for 10SrHA surface (1396.0 ± 35.6 kJ/mol/nm) in harmony with the previously calculated FN-surface interaction data (Fig. 2(a)). The FN never got fully detached from the 10SrHA surface (Fig.7(c)) and is attached to the surface via Val1 and Glu9 residues, the two most highly interacting residues (Fig.4(b)). For all other surfaces, FN got detached from the surface (Fig.7). The second highest value for the rupturing force was recorded for 30SrHA surface (1258.4 ± 58.5 kJ/mol/nm), followed by 0SrHA (1094.9 ± 78.7 kJ/mol/nm) and 20SrHA (979.2 ± 92.4 kJ/mol/nm). On the other hand, the unbinding time for all the surfaces was found to be beyond 200 ps (Table 3) with its highest value for 10SrHA surface (236.4 ± 8.8 ps).

**Table 3:** Rupture force and unbinding time, computed from Steered molecular dynamics simulation

| Code of Biomaterials surface | Sr Doping (mol%) | Rupture force (kJ/mol/nm) | Unbinding time (ps) |
| --- | --- | --- | --- |
| 0SrHA | 0 | $1094.9 \pm 78.7$ | $207.9 \pm 3.3$ |
| 10SrHA | 10 | $1396.0 \pm 35.6$ | $236.4 \pm 8.8$ |
| 20SrHA | 20 | $979.2 \pm 92.4$ | $208.7 \pm 53.6$ |
| 30SrHA | 30 | $1258.4 \pm 58.5$ | $220.9 \pm 27.5$ |

**Fig. 7:**
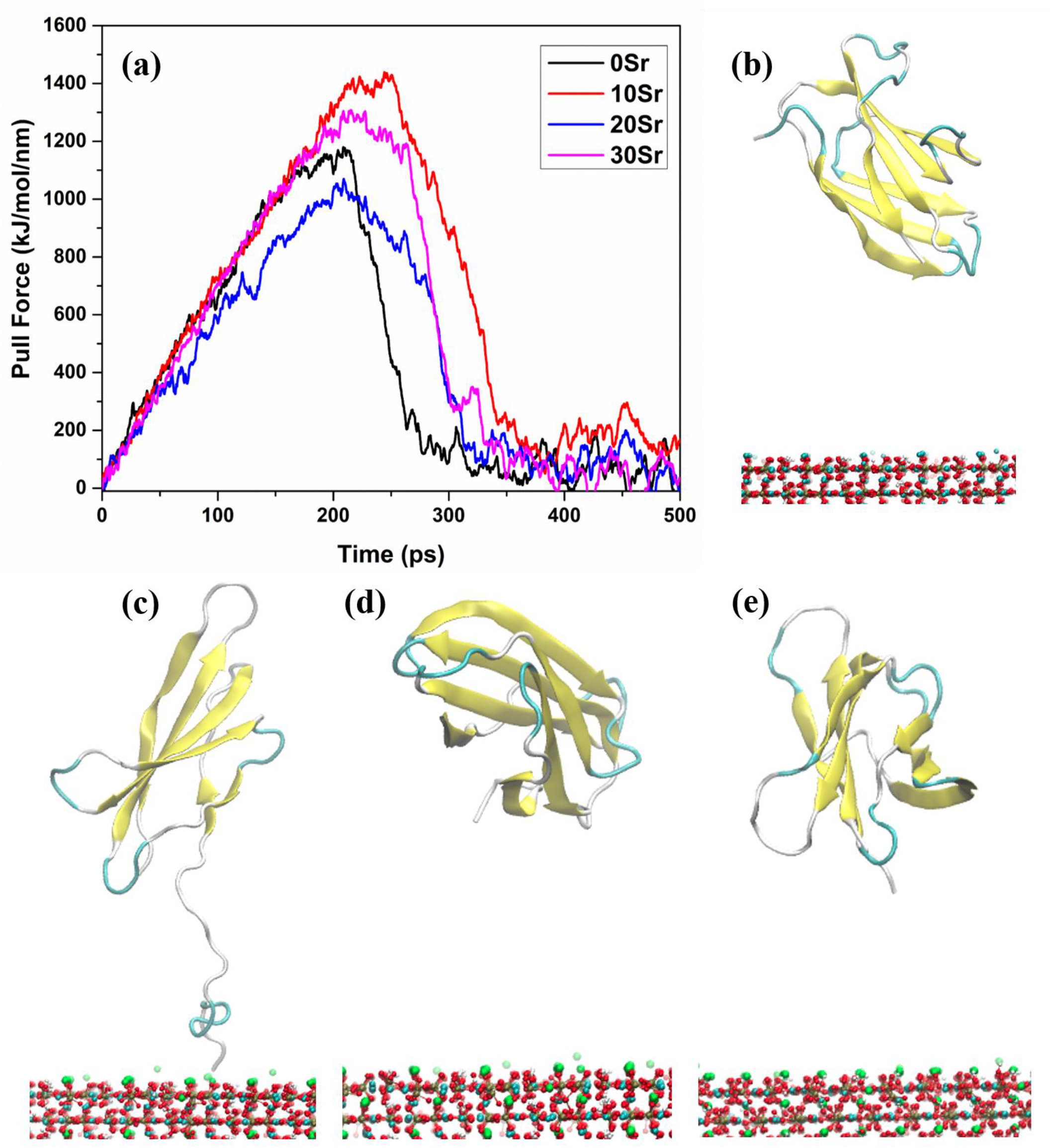
Sr^2+^ doping significantly regulates the binding affinity of FN. (a) Typical force curves for FN-undoped/Sr^2+^-doped HA system. Final configuration of the system after 500ps of SMD simulation for (b) 0SrHA, (c) 10SrHA, (d)20SrHA, and (e)30SrHA. Water molecules are not shown due to clarity. The secondary structure of FN has been represented with colour code: Yellow: β sheet, pale blue: turns, white: other residues. HA has been presented using CPK colouring scheme with atomic colour code: Red: O, pale blue: Ca, white: H, golden yellow: P, Green: Sr.

## 4. Discussion

In this section, the underlying causes of the recorded behaviour of the FN on undoped/Sr-doped HA surface will be analyzed. Moreover, a correlation among the changes in structural properties of FN module will also be drawn together with the different modes of interaction between protein and inorganic surface.

### 4.1 Influence of Sr^2+^ substitution on adsorption kinetics

It is now well-known that amino acids interact with the inorganic surface through electrostatic interaction and hydrogen bonds.^53, 54^ For HA, the electrostatic interaction occurs between the oppositely charged groups, i.e. Ca^2+^ from HA and COO^-^ of amino acids and PO_4_^3-^ of HA and NH_3_^+^ of the protein.^53^ Moreover, H-bond formation was also observed to happen between PO_4_^3-^ and NH_3_^+^ groups during the adsorption of BMP-7 on the HA (001) surface.^27^ The formation of strong Ca-O bonds was also noticed to mediate the interaction of HA and amino acids in its zwitterionic forms.^54^ Because of this reason, zwitterionic forms of many amino acids are prone to get adsorbed on the HA surface, as many studies have reported it previously.^55, 23, 24^ Quantum mechanical calculations reported an evidence of proton transfer from COOH^-^ group to HA surface during adsorption of Gly on HA (010) surface, resulting in the formation of Ca-O or O-Ca-O bond.^55^ Similar results were obtained for other amino acids, as well.^23, 24^

It is also evident that only specific residues of a protein interact with the HA surface during adsorption. Three specific binding sites were reported to be involved in protein-HA interaction.^56^

1. Phosphorylated amino acids
2. γ-carboxyglutamic acid of bone-Gla protein
3. Sequence of acidic amino acids like Asp or Glu rich sections of proteins

In the current study, it was found that, mainly the charged residues interacted with the inorganic surfaces by means of long-range coulombic interaction between oppositely charged groups of FN and HA. This interaction was found to be tailored by surface modification through Sr^2+^ doping.

The effect of Sr^2+^ doping can be understood in the following manner. Sr^2+^ is more reactive in nature, compared to Ca^2+^.^57^ The replacement of Ca^2+^ with Sr^2+^ causes a change in the nature of the non-bonded interaction between protein and surface. Because of this, additional structural/orientational changes of FN take place to compensate for the change in the environment. This structural rearrangement of protein, in turn, changes the number of interactive residues near the surface. Hence, the adsorbed state of FN becomes stronger or weaker. When there was no strontium (0SrHA), a few residues interacted with the surface (Table.2) and the FN module went through weak and strong adsorbed states, as seen in Fig 2(a) and 3(a). The structural stability was also disturbed, as seen from the high RMSD values (Fig.5(a)). The fluctuating number of contacts also supports this claim (Fig. 3(c)). When Sr^2+^ was introduced (10SrHA), the presence of more reactive Sr^2+^ ions brought more residues at the proximity of the surface, resulting in an increase in the attractive coulombic interaction energy, compared to 0SrHA (Table 2, Fig. 2(a)). The number of contacts and thus the number of H-bonds also increased. Also, FN remained in a relatively stable adsorbed state on the surface (Fig. 2(c) and 3(a)). The structural stability was also enhanced, as observed in Fig. 5(a).

When Sr^2+^ content was increased (20SrHA), the opposite effect was observed. Although the extra Sr^2+^ ions were located away from the FN module (Fig S3 in supporting information), the resulting repositioning of the local atoms due to their presence caused the movement of the Arg93, Val1, Ser2, and Thr94 away from the surface and Asp7 towards the surface (Fig S4 in supporting information). This resulted in a decrease in the overall coulombic interaction energy (Fig. 2(a)) while the interaction of Asp7 with the surface increased significantly (Fig.4). Although the electrostatic interaction energy decreased when compared to 10SrHA, it still remained higher than 0SrHA (Fig. 2(a)). Besides this, the number of contacts and H-bonds were also noticed to get decreased (Fig.2(c), 2(d) and 3(c)). Overall, these led to a relatively weaker adsorbed state of FN on 20SrHA surface, compared to 10SrHA. The condition slightly improved with the further increment of Sr content (30SrHA). In this case, Arg93 again moved to close proximity of the surface (Table 2), resulting in an increase in interaction energy, number of contacts, and H-bonds (Fig. 2 and 3(c)). But the presence of excess Sr^2+^ ions caused structural instability in FN, as seen from the high RMSD value (Fig. 5(a)). Despite this fact, all Sr-doped HA surfaces proved to better support FN adsorption, compared to the pure HA surface together with providing a favourable condition for cell-material interaction (Fig. 6).

### 4.2 Doping dependent unbinding of FN

Constant velocity SMD simulation was performed on the system to mimic the AFM experiments. Here, a force was applied on the FN and was pulled away from the surface together with the calculation of pulling force. A correlation can be drawn among the rupture force, the structural changes of the FN module and the interaction between the surface and protein. The highest rupture force was obtained for 10SrHA, because of the highly attractive interaction between FN and surface (Fig. 7(a) and 2(a)). The interaction was so strong that the FN did not detach from the surface at the end of 500 ps (Fig.7(c)). Because of the high RMSD (and less interaction), the rupture force was found to be less in 0SrHA and 30SrHA (Fig. 7(a) and Table 3). One notable finding is that the rupture force was noticed to be least in 20SrHA than pure HA, although the average coulombic interaction was higher for 20SrHA (Table 1). Repulsive non-bonded interaction and a smaller number of interacting residues may be the probable reasons behind the obtained trend (Fig. 2(b) and 3(c). However, the comparison between the coulombic interaction between 20SrHA and 0SrHA indicates that, the FN was strongly adsorbed on 20SrHA like its undoped counterpart.

## 5. Conclusions

In the current work, we have explored the influences of doping of biologically relevant metal ions like Sr^2+^ in apatite structure on protein adsorption mechanism. Following are the key conclusions drawn from the present study.

a. Adsorption of FN can be tailored by modulating the amount of Sr^2+^ ions inside HA crystal structure. In general, strontium doped HA supports the adsorption of fibronectin better than pure HA surface. FN exhibited the highest affinity towards 10 mol% Sr doped HA.
b. Sr^2+^ ions modify the adsorption kinetics by means of altering the number of interacting residues, which result from the rearrangement of the positions of the atoms of the protein, in an attempt to compensate for the effects of surface modification.
c. Secondary structure of the FN was mildly affected by Sr^2+^ doping. A slight change in the percentage of the β-sheet was observed with an increase of Sr^2+^ content. Besides this, non-monotonous behaviour of RMSD and RMSF was observed. This can be attributed to the structural changes of protein induced by Sr^2+^ doping in apatite structure.
d. RGD motif was exposed to the solvent site irrespective of the nature of the biomaterials surface. It indicates that, the cell-material interaction will not be compromised with Sr^2+^ doping.

Summarizing, we explored the effects of doping of one biologically relevant metal ion, Sr^2+^, on fibronectin (FN) adsorption on HA (001) surface. The presented results can rationalize the experimentally proven better cytocompatibility of Sr-doped HA.^35^ In this article, we have only considered the effects of Sr^2+^ ions substituted in the Ca(1) positions of HA. A future study can be designed to explore the physicochemical effects of metal ion substitution at different lattice sites on adsorption kinetics, which can pave a pathway to design the next generation biometerials.

## Supporting information

Contains figures not included in the main manuscript

## Acknowledgments

The authors acknowledge the Thematic Unit of Excellence on Computational Materials Science (TUE-CMS) of Solid State and Structural Chemistry Unit (SSCU) at IISc, Bangalore, for providing access to the high-performance computing facilities. BB acknowledges IMPRINT project funded by DST, SERB, Govt. of India for financial assistance as well as Abdul Kalam Technology Innovation National Fellowship, funded by DST-INAE, Govt. of India. SB and PKM acknowledge support through SERB, IRHPA through IPA/2020/000034.

## References

(1) Basu, B. Biomaterials for Musculoskeletal Regeneration; Springer, 2017.

(2) Basu, B. Biomaterials Science and Tissue Engineering: Principles and Methods; Cambridge University Press, 2017.

(3) Basu, B.; Katti, D. S.; Kumar, A. Advanced Biomaterials: Fundamentals, Processing, and Applications; John Wiley & Sons, 2010.

(4) Gibbons, R. J.; Hay, D. I.; Childs Iii, W. C.; Davis, G. Role of Cryptic Receptors (Cryptitopes) in Bacterial Adhesion to Oral Surfaces. Arch. Oral Biol. 1990, 35, S107–S114.

(5) Ozboyaci, M.; Kokh, D. B.; Corni, S.; Wade, R. C. Modeling and Simulation of Protein–Surface Interactions: Achievements and Challenges. Q. Rev. Biophys. 2016, 49.

(6) Dufrêne, Y. F.; Marchal, T. G.; Rouxhet, P. G. Influence of Substratum Surface Properties on the Organization of Adsorbed Collagen Films: In Situ Characterization by Atomic Force Microscopy. Langmuir 1999, 15 (8), 2871–2878.

(7) Bergkvist, M.; Carlsson, J.; Oscarsson, S. Surface-dependent Conformations of Human Plasma Fibronectin Adsorbed to Silica, Mica, and Hydrophobic Surfaces, Studied with Use of Atomic Force Microscopy. J. Biomed. Mater. Res. Part A An Off. J. Soc. Biomater. Japanese Soc. Biomater. Aust. Soc. Biomater. Korean Soc. Biomater. 2003, 64 (2), 349–356.

(8) Henry, M.; Dupont-Gillain, C.; Bertrand, P. Conformation Change of Albumin Adsorbed on Polycarbonate Membranes as Revealed by ToF-SIMS. Langmuir 2003, 19 (15), 6271–6276.

(9) Hagiwara, T.; Sakiyama, T.; Watanabe, H. Molecular Simulation of Bovine β-Lactoglobulin Adsorbed onto a Positively Charged Solid Surface. Langmuir 2009, 25 (1), 226–234.

(10) Hoefling, M.; Monti, S.; Corni, S.; Gottschalk, K. E. Interaction of β-Sheet Folds with a Gold Surface. PLoS One 2011, 6 (6), e20925.

(11) Iori, F.; Corni, S.; Di Felice, R. Unraveling the Interaction between Histidine Side Chain and the Au (111) Surface: A DFT Study. J. Phys. Chem. C 2008, 112 (35), 13540–13545.

(12) Mulheran, P. A.; Connell, D. J.; Kubiak-Ossowska, K. Steering Protein Adsorption at Charged Surfaces: Electric Fields and Ionic Screening. RSC Adv. 2016, 6 (77), 73709–73716.

(13) Tokarczyk, K.; Kubiak-Ossowska, K.; Jachimska, B.; Mulheran, P. A. Energy Landscape of Negatively Charged BSA Adsorbed on a Negatively Charged Silica Surface. J. Phys. Chem. B 2018, 122 (14), 3744–3753.

(14) Patwardhan, S. V; Emami, F. S.; Berry, R. J.; Jones, S. E.; Naik, R. R.; Deschaume, O.; Heinz, H.; Perry, C. C. Chemistry of Aqueous Silica Nanoparticle Surfaces and the Mechanism of Selective Peptide Adsorption. J. Am. Chem. Soc. 2012, 134 (14), 6244–6256.

(15) Tosaka, R.; Yamamoto, H.; Ohdomari, I.; Watanabe, T. Adsorption Mechanism of Ribosomal Protein L2 onto a Silica Surface: A Molecular Dynamics Simulation Study. Langmuir 2010, 26 (12), 9950–9955.

(16) Lecot, S.; Chevolot, Y.; Phaner-Goutorbe, M.; Yeromonahos, C. Impact of Silane Monolayers on the Adsorption of Streptavidin on Silica and Its Subsequent Interactions with Biotin: Molecular Dynamics and Steered Molecular Dynamics Simulations. J. Phys. Chem. B 2020, 124 (31), 6786–6796.

(17) Utesch, T.; Daminelli, G.; Mroginski, M. A. Molecular Dynamics Simulations of the Adsorption of Bone Morphogenetic Protein-2 on Surfaces with Medical Relevance. Langmuir 2011, 27 (21), 13144–13153.

(18) Mücksch, C.; Urbassek, H. M. Accelerated Molecular Dynamics Study of the Effects of Surface Hydrophilicity on Protein Adsorption. Langmuir 2016, 32 (36), 9156–9162.

(19) Mazouz, Z.; Mokni, M.; Fourati, N.; Zerrouki, C.; Barbault, F.; Seydou, M.; Kalfat, R.; Yaakoubi, N.; Omezzine, A.; Bouslema, A. Computational Approach and Electrochemical Measurements for Protein Detection with MIP-Based Sensor. Biosens. Bioelectron. 2020, 151, 111978.

(20) Xie, Y.; Liu, M.; Zhou, J. Molecular Dynamics Simulations of Peptide Adsorption on Self-Assembled Monolayers. Appl. Surf. Sci. 2012, 258 (20), 8153–8159.

(21) Xie, Y.; Liao, C.; Zhou, J. Effects of External Electric Fields on Lysozyme Adsorption by Molecular Dynamics Simulations. Biophys. Chem. 2013, 179, 26–34.

(22) Basu, S.; Basu, B. Doped Biphasic Calcium Phosphate: Synthesis and Structure. J. Asian Ceram. Soc. 2019, 7 (3), 265–283.

(23) Lou, Z.; Zeng, Q.; Chu, X.; Yang, F.; He, D.; Yang, M.; Xiang, M.; Zhang, X.; Fan, H. First-Principles Study of the Adsorption of Lysine on Hydroxyapatite (1 0 0) Surface. Appl. Surf. Sci. 2012, 258 (11), 4911–4916.

(24) Almora-Barrios, N.; Austen, K. F.; de Leeuw, N. H. Density Functional Theory Study of the Binding of Glycine, Proline, and Hydroxyproline to the Hydroxyapatite (0001) and (0110) Surfaces. Langmuir 2009, 25 (9), 5018–5025.

(25) Almora-Barrios, N.; de Leeuw, N. H. A Density Functional Theory Study of the Interaction of Collagen Peptides with Hydroxyapatite Surfaces. Langmuir 2010, 26 (18), 14535–14542.

(26) Dong, X.-L.; Qi, W.; Tao, W.; Ma, L.-Y.; Fu, C.-X. The Dynamic Behaviours of Protein BMP-2 on Hydroxyapatite Nanoparticles. Mol. Simul. 2011, 37 (13), 1097–1104.

(27) Zhou, H.; Wu, T.; Dong, X.; Wang, Q.; Shen, J. Adsorption Mechanism of BMP-7 on Hydroxyapatite (001) Surfaces. Biochem. Biophys. Res. Commun. 2007, 361 (1), 91–96.

(28) Gu, H.; Xue, Z.; Wang, M.; Yang, M.; Wang, K.; Xu, D. Effect of Hydroxyapatite Surface on BMP-2 Biological Properties by Docking and Molecular Simulation Approaches. J. Phys. Chem. B 2019, 123 (15), 3372–3382.

(29) Shen, J. W.; Wu, T.; Wang, Q.; Pan, H. H. Molecular Simulation of Protein Adsorption and Desorption on Hydroxyapatite Surfaces. Biomaterials 2008, 29 (5), 513–532.

(30) Liao, C.; Xie, Y.; Zhou, J. Computer Simulations of Fibronectin Adsorption on Hydroxyapatite Surfaces. RSC Adv. 2014, 4 (30), 15759–15769.

(31) Basu, S.; Gorai, B.; Basu, B.; Maiti, P. K. Electric Field-Mediated Fibronectin– Hydroxyapatite Interaction: A Molecular Insight. J. Phys. Chem. B 2021, 125 (1), 3–16.

(32) Pankov, R.; Yamada, K. M. Fibronectin at a Glance. J. Cell Sci. 2002, 115 (20), 3861–3863.

(33) Šupová, M. Substituted Hydroxyapatites for Biomedical Applications: A Review. Ceram. Int. 2015, 41 (8), 9203–9231.

(34) Marie, P. J.; Ammann, P.; Boivin, G.; Rey, C. Mechanisms of Action and Therapeutic Potential of Strontium in Bone. Calcif. Tissue Int. 2001, 69 (3), 121–129.

(35) Capuccini, C.; Torricelli, P.; Boanini, E.; Gazzano, M.; Giardino, R.; Bigi, A. Interaction of Sr-doped Hydroxyapatite Nanocrystals with Osteoclast and Osteoblast-like Cells. J. Biomed. Mater. Res. Part A An Off. J. Soc. Biomater. Japanese Soc. Biomater. Aust. Soc. Biomater. Korean Soc. Biomater. 2009, 89 (3), 594–600.

(36) Saidak, Z.; Marie, P. J. Strontium Signaling: Molecular Mechanisms and Therapeutic Implications in Osteoporosis. Pharmacol. Ther. 2012, 136 (2), 216–226.

(37) Zhou, J.; Han, Y.; Lu, S. Direct Role of Interrod Spacing in Mediating Cell Adhesion on Sr-HA Nanorod-Patterned Coatings. Int. J. Nanomedicine 2014, 9, 1243.

(38) Xu, Y.; An, L.; Chen, L.; Xu, H.; Zeng, D.; Wang, G. Controlled Hydrothermal Synthesis of Strontium-Substituted Hydroxyapatite Nanorods and Their Application as a Drug Carrier for Proteins. Adv. Powder Technol. 2018, 29 (4), 1042–1048.

(39) Huang, B.; He, M.; Zhang, K.; Sun, S.; Lin, Z.; Chen, D.; Li, T.; Chen, X. Rotation-Induced Secondary Structure Losses and Bioactivity Changes of Bone Morphogenetic Protein-2 on Strontium-Substituted Hydroxyapatite Surfaces. Appl. Surf. Sci. 2020, 511, 145623.

(40) Liu, Q.; Xue, Z.; Xu, D. Molecular Dynamics Characterization of Sr-Doped Biomimetic Hydroxyapatite Nanoparticles. J. Phys. Chem. C 2020, 124 (36), 19704–19715.

(41) Main, A. L.; Harvey, T. S.; Baron, M.; Boyd, J.; Campbell, I. D. The Three-Dimensional Structure of the Tenth Type III Module of Fibronectin: An Insight into RGD-Mediated Interactions. Cell 1992, 71 (4), 671–678.

(42) Liao, C.; Xie, Y.; Zhou, J. Computer Simulations of Fibronectin Adsorption on Hydroxyapatite Surfaces. RSC Adv. 2014, 4 (30), 15759–15769.

(43) Basu, S.; Ghosh, A.; Barui, A.; Basu, B. (Fe/Sr) Codoped Biphasic Calcium Phosphate with Tailored Osteoblast Cell Functionality. ACS Biomater. Sci. Eng. 2018, 4 (3), 857–871.

(44) Lin, T. J.; Heinz, H. Accurate Force Field Parameters and PH Resolved Surface Models for Hydroxyapatite to Understand Structure, Mechanics, Hydration, and Biological Interfaces. J. Phys. Chem. C 2016, 120 (9), 4975–4992.

(45) Corno, M.; Rimola, A.; Bolis, V.; Ugliengo, P. Hydroxyapatite as a Key Biomaterial: Quantum-Mechanical Simulation of Its Surfaces in Interaction with Biomolecules. Phys. Chem. Chem. Phys. 2010, 12 (24), 6309–6329.

(46) Bussi, G.; Donadio, D.; Parrinello, M. Canonical Sampling through Velocity Rescaling. J. Chem. Phys. 2007, 126 (1), 14101.

(47) Parrinello, M.; Rahman, A. Polymorphic Transitions in Single Crystals: A New Molecular Dynamics Method. J. Appl. Phys. 1981, 52 (12), 7182–7190.

(48) Hess, B.; Bekker, H.; Berendsen, H. J. C.; Fraaije, J. G. E. M. LINCS: A Linear Constraint Solver for Molecular Simulations. J. Comput. Chem. 1997, 18 (12), 1463–1472.

(49) Darden, T.; York, D.; Pedersen, L. Particle Mesh Ewald: An N Log (N) Method for Ewald Sums in Large Systems. J. Chem. Phys. 1993, 98 (12), 10089–10092.

(50) Abraham, M. J.; Murtola, T.; Schulz, R.; Páll, S.; Smith, J. C.; Hess, B.; Lindahl, E. GROMACS: High Performance Molecular Simulations through Multi-Level Parallelism from Laptops to Supercomputers. SoftwareX 2015, 1, 19–25.

(51) Lindorff-Larsen, K.; Piana, S.; Palmo, K.; Maragakis, P.; Klepeis, J. L.; Dror, R. O.; Shaw, D. E. Improved Side-chain Torsion Potentials for the Amber Ff99SB Protein Force Field. Proteins Struct. Funct. Bioinforma. 2010, 78 (8), 1950–1958.

(52) Humphrey, W.; Dalke, A.; Schulten, K. VMD: Visual Molecular Dynamics. J. Mol. Graph. 1996, 14 (1), 33–38.

(53) Wang, Q.; Wang, M.; Wang, K.; Liu, Y.; Zhang, H.; Lu, X.; Zhang, X. Computer Simulation of Biomolecule–Biomaterial Interactions at Surfaces and Interfaces. Biomed. Mater. 2015, 10 (3), 32001.

(54) Basu, S.; Basu, B. Unravelling Doped Biphasic Calcium Phosphate: Synthesis to Application. ACS Appl. Bio Mater. 2019, 2 (12), 5263–5297.

(55) Jimenez-Izal, E.; Chiatti, F.; Corno, M.; Rimola, A.; Ugliengo, P. Glycine Adsorption at Nonstoichiometric (010) Hydroxyapatite Surfaces: A B3LYP Study. J. Phys. Chem. C 2012, 116 (27), 14561–14567.

(56) Fujisawa, R.; Wada, Y.; Nodasaka, Y.; Kuboki, Y. Acidic Amino Acid-Rich Sequences as Binding Sites of Osteonectin to Hydroxyapatite Crystals. Biochim. Biophys. Acta (BBA)-Protein Struct. Mol. Enzymol. 1996, 1292 (1), 53–60.

(57) Greenwood, N. N.; Earnshaw, A. Chemistry of the Elements; Elsevier, 2012.

