## Supplementary material for "Computational study on Strontium ion modified Fibronectin-Hydroxyapatite interaction": Contains figures not included in the main manuscript

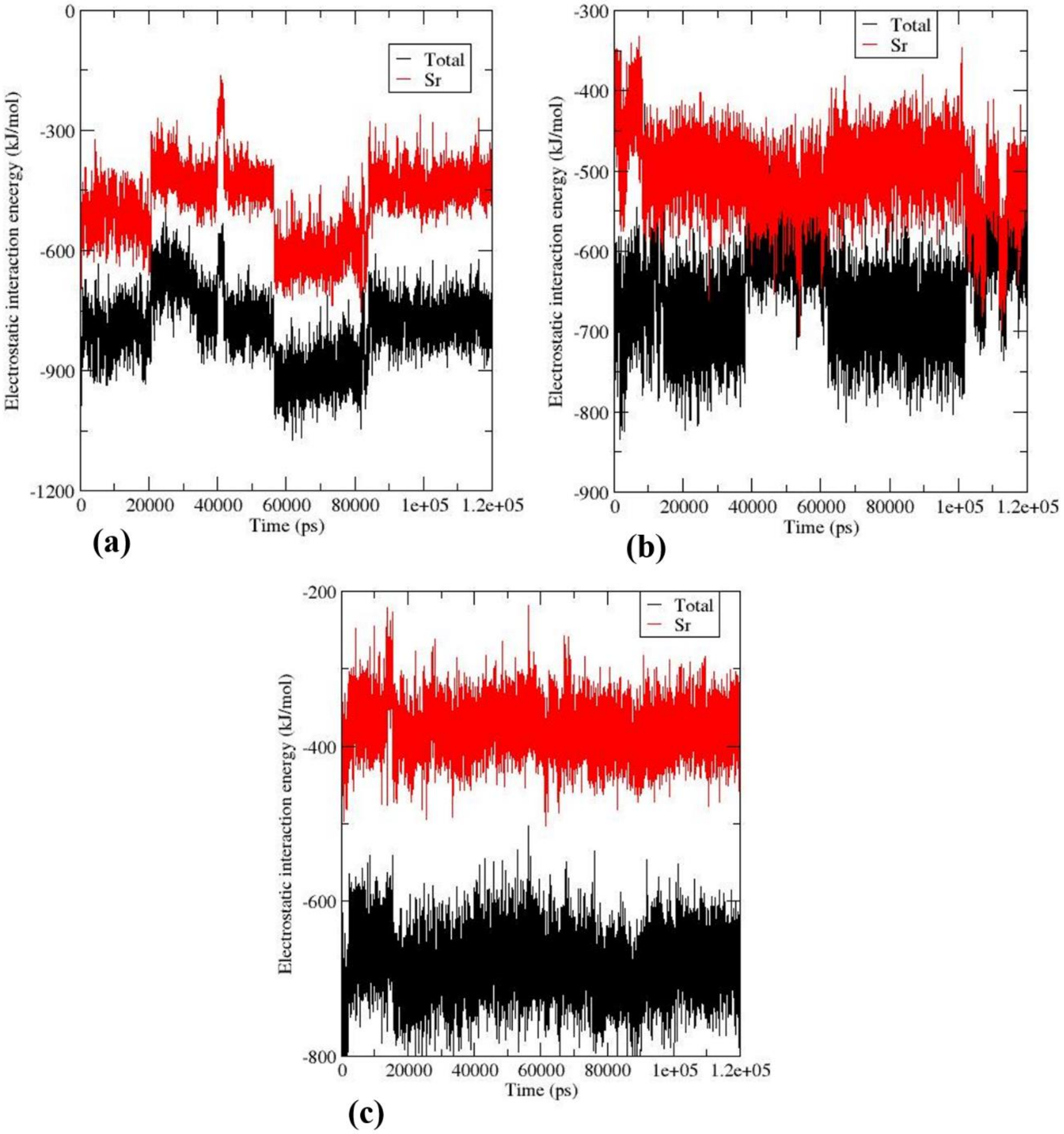


**Fig S1:** Comparison of total electrostatic interaction between FN and surface with that between $Sr^{2+}$ and FN for (a)10SrHA, (b)20SrHA, and (c) 30SrHA.

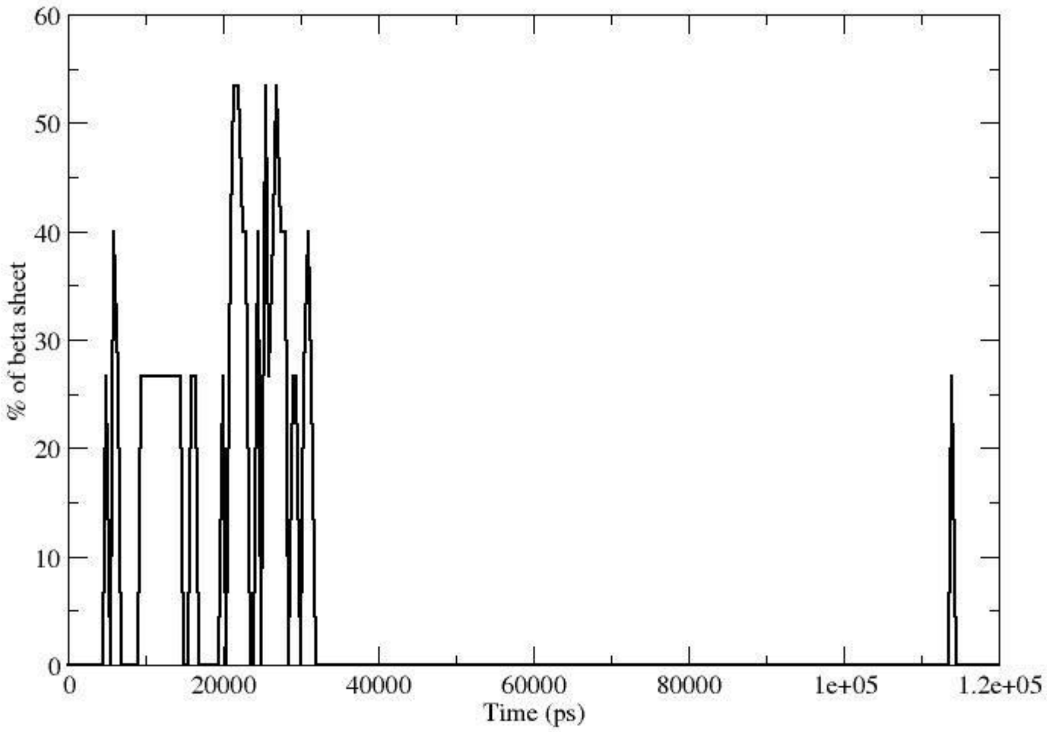


**Fig S2:** Temporal evolution of percentage of β-sheet in ‘area2’ (residue74-88) of FN.

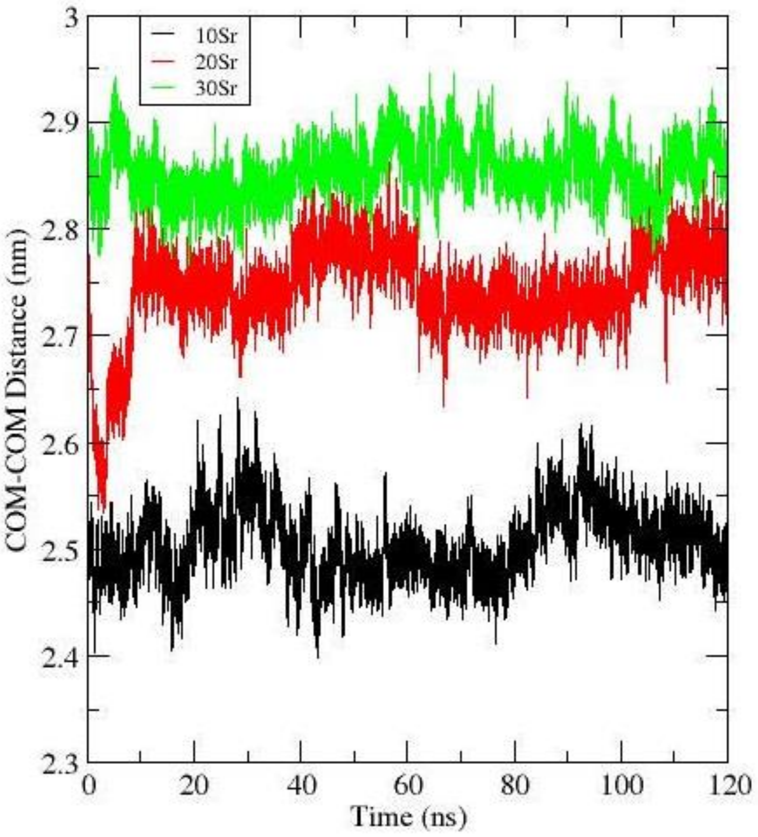


**Fig S3:** COM-COM distance between FN and $Sr^{2+}$ ions. Increasing distance indicates $Sr^{2+}$ ions were substituted away from FN module.

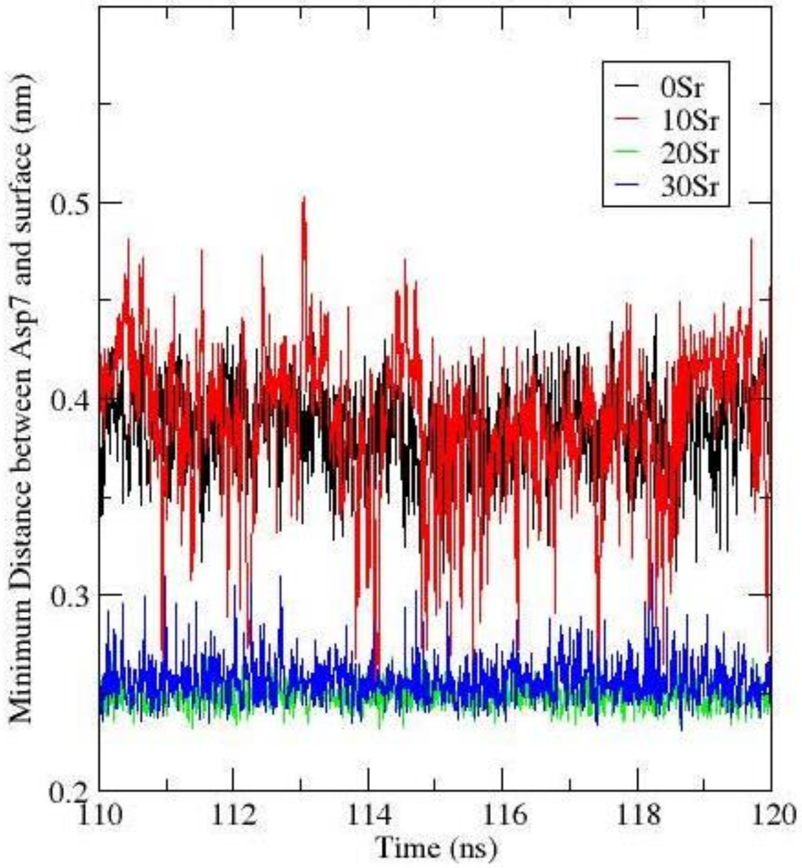


**Fig S4:** Minimum distance between Asp7 residue and undoped/Sr-doped HA surfaces.

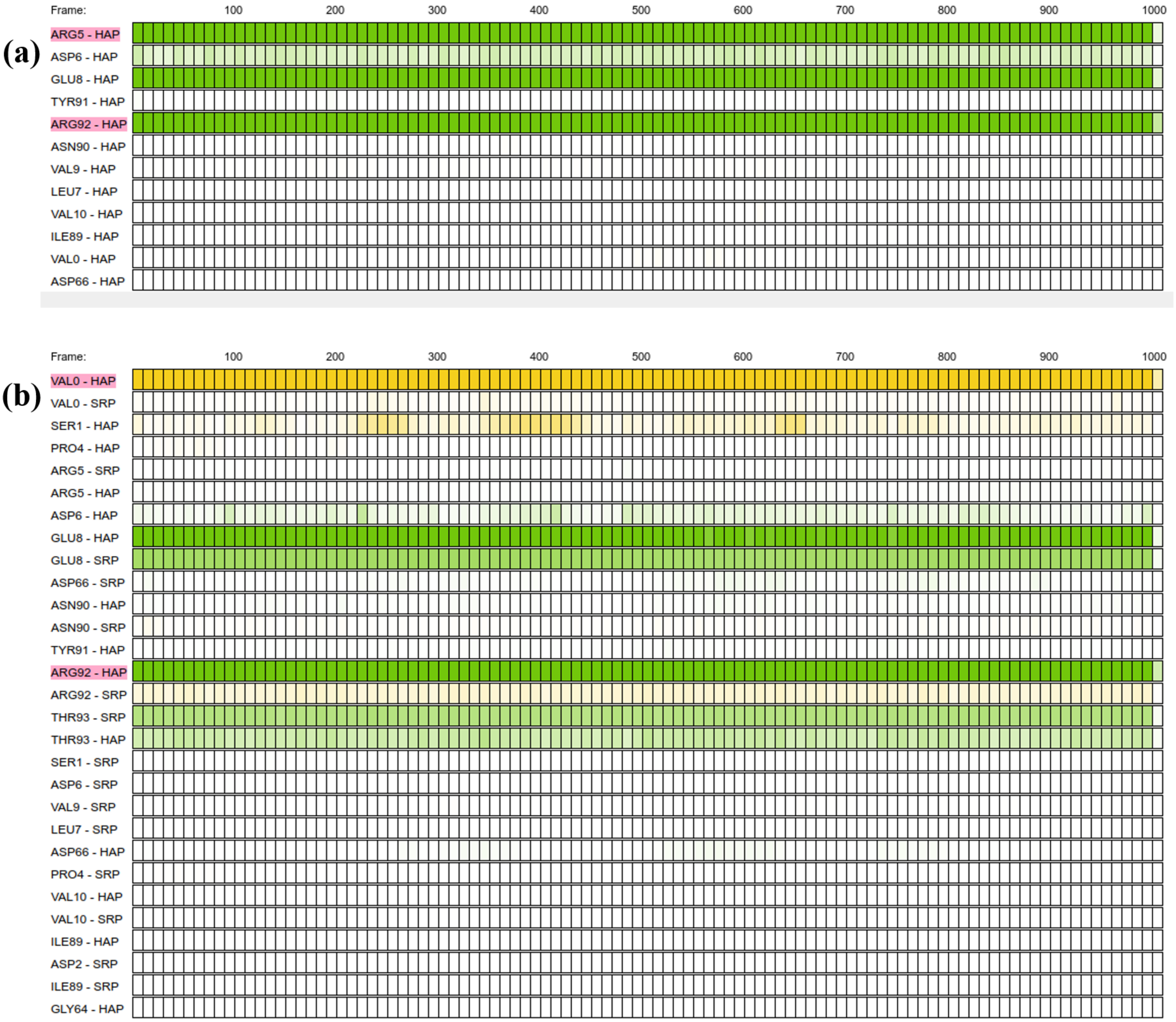
(a)
Frame:
100
200
300
400
500
600
700
800
900
1000
ARG5 - HAP
ASP6 - HAP
GLU8 - HAP
TYR91 - HAP
ARG92 - HAP
ASN90 - HAP
VAL9 - HAP
LEU7 - HAP
VAL10 - HAP
ILE89 - HAP
VAL0 - HAP
ASP66 - HAP
(b)
Frame:
100
200
300
400
500
600
700
800
900
1000
VAL0 - HAP
VAL0 - SRP
SER1 - HAP
PRO4 - HAP
ARG5 - SRP
ARG5 - HAP
ASP6 - HAP
GLU8 - HAP
GLU8 - SRP
ASP66 - SRP
ASN90 - HAP
ASN90 - SRP
TYR91 - HAP
ARG92 - HAP
ARG92 - SRP
THR93 - SRP
THR93 - HAP
SER1 - SRP
ASP6 - SRP
VAL9 - SRP
LEU7 - SRP
ASP66 - HAP
PRO4 - SRP
VAL10 - HAP
VAL10 - SRP
ILE89 - HAP
ASP2 - SRP
ILE89 - SRP
GLY64 - HAP

(c)

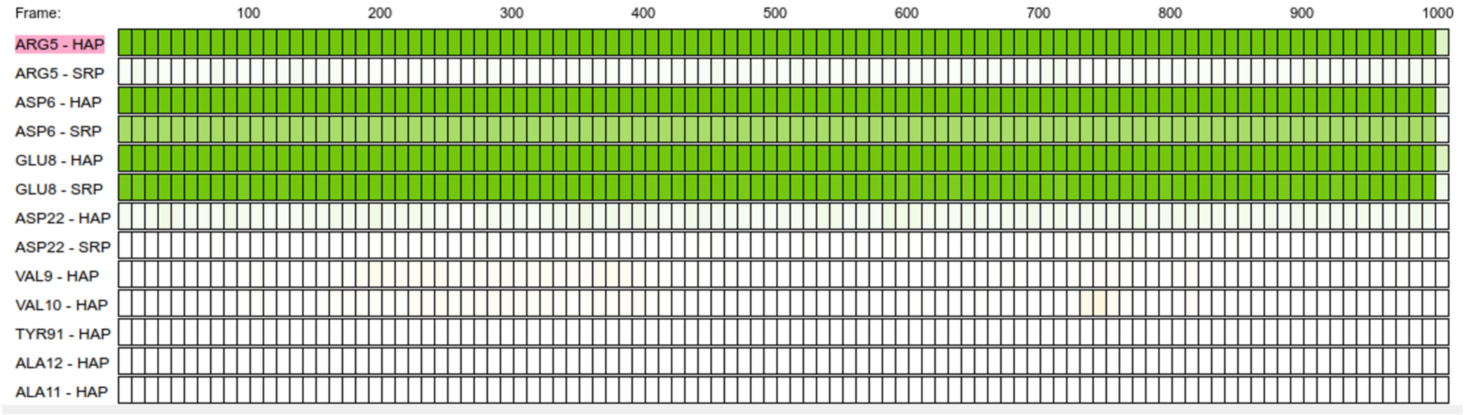


(d)

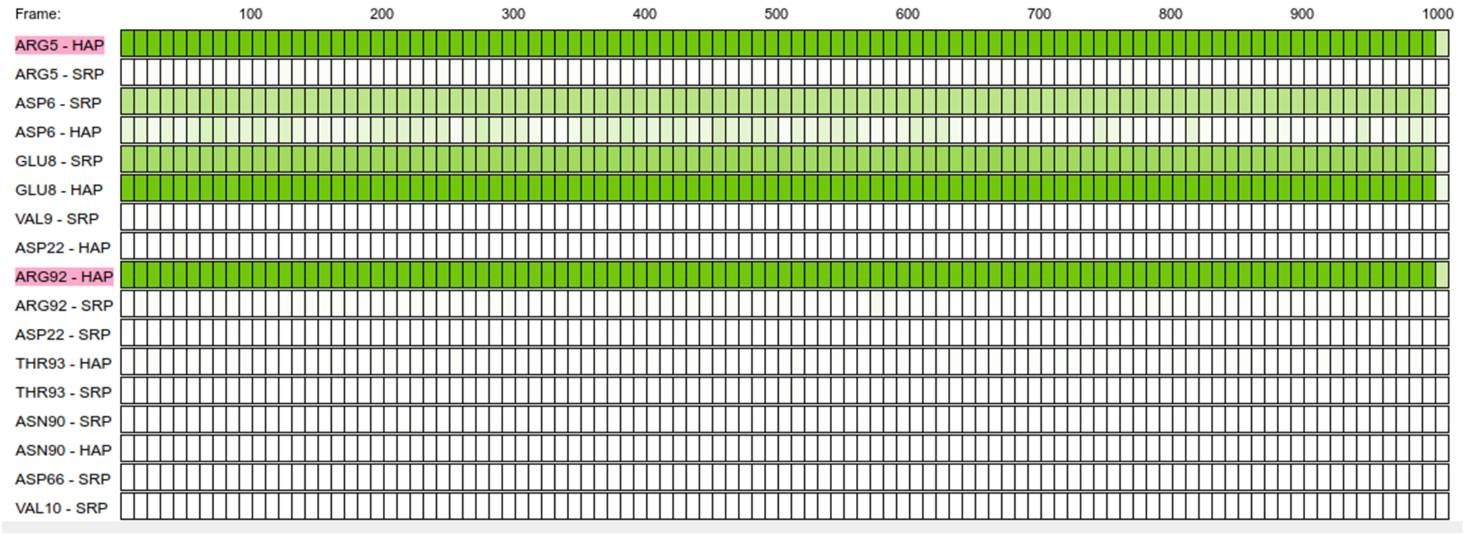


**Fig S5:** Timeline of the contact analysis of the last 1000 frames (110-120 ns) of the simulation for (a)0SrHA, (b) 10SrHA, (c) 20SrHA, and (d) 30SrHA, obtained using PyContact tool.[1] Pink colour at the left most panels highlights hydrogen bond forming residues. Green and yellow colour indicates side chain-side chain and side chain-backbone interactions, respectively. Strength of the interaction is proportional to the colour intensity. Cut off distance for the analysis was 5Å.

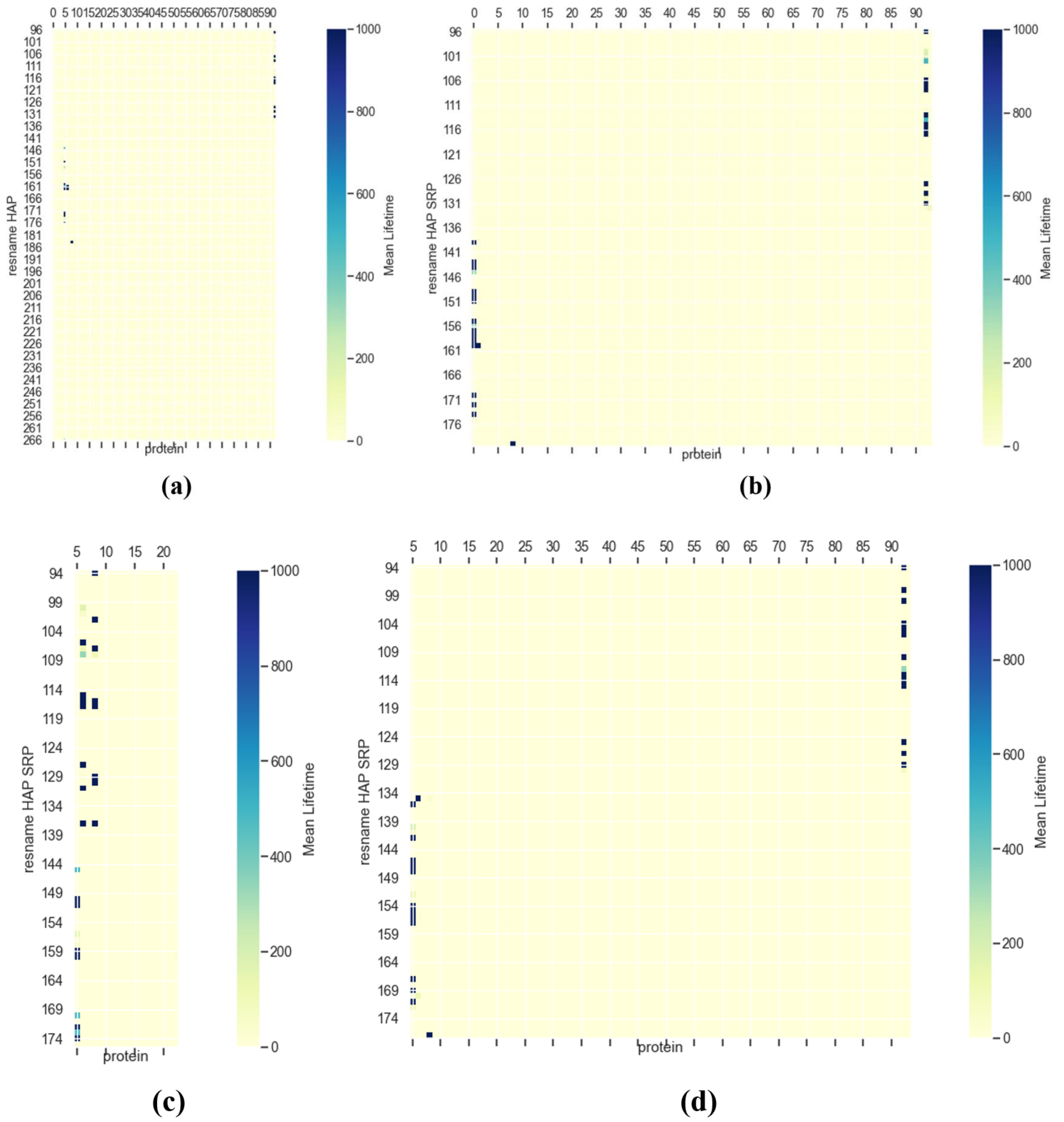


**Fig S6:** FN-materials surface Contact map for (a)0SrHA, (b) 10SrHA, (c) 20SrHA, and (d) 30SrHA, obtained using PyContact tool.[1] A contact has been formed when two atoms, one from FN and other from the surface are within a distance of 5Å. The mean life time (a.u) of the contacts formed between residues of FN and surface is estimated and shown.

### References

(1) Scheurer, M.; Rodenkirch, P.; Siggel, M.; Bernardi, R. C.; Schulten, K.; Tajkhorshid, E.; Rudack, T. PyContact: Rapid, Customizable, and Visual Analysis of Noncovalent Interactions in MD Simulations. *Biophys. J.* **2018**, *114* (3), 577–583.
